# Dorsal hippocampus represents locations to avoid as well as locations to approach during approach-avoidance conflict

**DOI:** 10.1101/2024.03.10.584295

**Authors:** Olivia L. Calvin, Matthew T. Erickson, Cody J. Walters, A. David Redish

## Abstract

Worrying about perceived threats is a hallmark of multiple psychological disorders including anxiety. This concern about future events is particularly important when an individual is faced with an approach-avoidance conflict. Potential goals to approach are known to be represented in the dorsal hippocampus during theta sweeps. Similarly, important non-local information is represented during hippocampal high synchrony events (HSEs), which are correlated with sharp-wave ripples (SWRs). It is likely that potential future threats may be similarly represented. We examined how threats and rewards were represented within the hippocampus during approach-avoidance conflicts in rats faced with a predator-like robot guarding a food reward. We found representations of the pseudo-predator during HSEs when hesitating in the nest, and during theta prior to retreating as the rats approached the pseudo-predator. After the first attack, we observed new place fields appearing at the location of the robot (not the location the rat was when attacked). The anxiolytic diazepam reduced anxiety-like behavior and altered hippocampal local field potentials, including reducing SWRs, suggesting that one potential mechanism of diazepam’s actions may be through altered representations of imagined threat. These results suggest that hippocampal representation of potential threats could be an important mechanism that underlies worry and a potential target for anxiolytics.

## Introduction

Theories have long hypothesized that worry and anxiety involves imagination, specifically the ability to mentally simulate negative future outcomes. Indeed, the psychiatry literature has largely assumed this to be the case, positing that anxiety disorders result from cognitive schemas that distort one’s expectations and evaluations of the future [1–5]. Much theorizing has revolved around the notion of episodic future thinking in planning and goal-directed decision-making [6–8], but primarily in conditions of conflict between multiple positive outcomes (approach-approach conflict) [9,10]. In contrast, anxiety is thought to arise from conflict between motivations towards taking actions that one wants to both approach and avoid (approach-avoidance conflict) [11–14]. While theoretical work has long suggested a role in future thinking in these anxiety-inducing situations [4,5,15–17], and a specific role for planning-related structures such as hippocampus and prefrontal cortex therein [18], neurophysiological studies have concentrated on the emotional dimension of anxiety while largely ignoring the role of prospection [17,19–23].

Studies building on the spatial navigation literature have revealed that the hippocampus encodes mental simulations of hypothetical episodic scenarios during planning, sweeping paths forward in both space and time to the next goal or subgoal [24–29]. Similarly, studies building on the same spatial navigation literature have revealed that the hippocampus replays important information during behavioral pauses, including both appetitive [30–35] and aversive [36] memory recall. However, it remains unclear what role, if any, hippocampal fictive representations play in threat-based approach-avoidance conflict.

One such task is the predator-inhabited foraging task, an approach-avoidance conflict task that requires a rat to forage for food in the presence of a hostile robotic predator designed to mimic the hazardous environmental conditions faced by rodents in the wild [37]. The predator-inhabited foraging task (colloquially known as the “robogator” task) elicits a variety of anxiety-like behaviors in rats [37–39], including choice-point hesitation at the nest entrance, which is similar to the stretch-attend posture, thought to arise from risk-assessment processes [40–43], mid-track retreat decisions, which are likely a kind of change-of-mind event [38,39], and slowing down during foraging, likely due to nervous concern about imminent dangers [38,44].

Consistent with this hypothesis, these behaviors are modulated by anti-anxiety drugs, particularly diazepam [39]. These behaviors depend on a complex neural circuit, including the amygdala [37,38], the bed nucleus of the stria terminalis [44], and the periaqueductal gray [45], as well as hippocampus [46]. Given the well-established role for hippocampus in prospective imagination of targets to approach, it becomes a particularly interesting question of how hippocampal representations encode the conflicting aspects of futures to approach and futures to avoid, and a particularly interesting question of how those drugs affect hippocampal processes.

The hippocampus shows two distinct states, identifiable by the oscillatory regimes revealed by the local field potential (LFP): a theta state, in which the local field potential shows prominent 6-10 Hz (theta) and 15-25 Hz (beta) oscillations and a non-theta state (termed large-amplitude irregular activity or LIA) in which the local field potential is dominated by 1-4 Hz delta band oscillations, punctuated by transient 150-250 Hz 150 ms events called sharp-wave ripples (SWRs) [47–52].

Hippocampal pyramidal cells are well-known to encode contextual and spatial information key to episodic events within a task, particularly through the well-studied place fields that represent locations in spatial environments [47,48]. Each theta cycle entails a descending phase (as recorded from the hippocampal layer) in which pyramidal cell firing reflects the current location of the animal, and an ascending phase in which pyramidal cell firing reflects the potential future paths of the animal [24,25,27,53–55]. On an avoidance task, transient representations of the danger zone have been identified during the non-local phase of theta [56]. However, it remains unknown how theta sweeps reflect that conflict between approach and avoidance.

While in LIA states, the hippocampal pyramidal cell population bursts in transient high synchrony events (HSEs) that reflect important information about the environment [29,31–35]. The specifics of the extent to which these transient events reflect planning [29,31,35,57], memory [31,32,35,58,59], learning [60–64] or some other process, such as development or maintenance of the spatial map [33,65–67] remains unknown. However, they have been observed to encode negative locations (shock-delivery locations) after novel experiences [36] and have been suggested to be keys to anxiety [68].

We sought to bring the anxiety, prospection, and ethology literatures together in order to interrogate the neural basis of negatively-valenced episodic thinking. To do this, we recorded neural activity from dorsal hippocampal ensembles on an approach-avoidance task in which rats ran between food locations, one of which was guarded by a robot pseudo-predator. We found hippocampal representational changes that created both sequential and discrete non-local hippocampal representations that co-occured with anxiety-like behaviors. We then found that diazepam had strong effects on exactly those hippocampal processes that contained anxiety-related information during those anxiety-like behaviors. The findings reported below suggest that the dorsal hippocampus conveys both reward-related and threat-related information in approach-avoidance conflict environments, and further suggest a role for the hippocampus in negatively-valenced imagination, thus providing a novel neural mechanism that could underlie a long-hypothesized psychological key to anxiety.

## Results

Neural ensembles were recorded from silicon probes implanted bilaterally into CA1 from rats (n=6, 3M 3F) facing an approach-avoidance conflict in which food rewards at one end of a linear track were sometimes interrupted by the attack of a robot pseudo-predator. This conflict is similar to earlier versions of the robogator task [37,38,44], but increased the number of laps run in each session and required the rat to run past the robot to receive the food reward (Figure 1A), which allowed us to measure the spatial representations of near-robot locations. Rats ran one 1 hr session each day. Rats first learned to run to alternate ends of a linear track for small (45mg) food rewards that would supply their daily nutrition.

**Figure 1.**
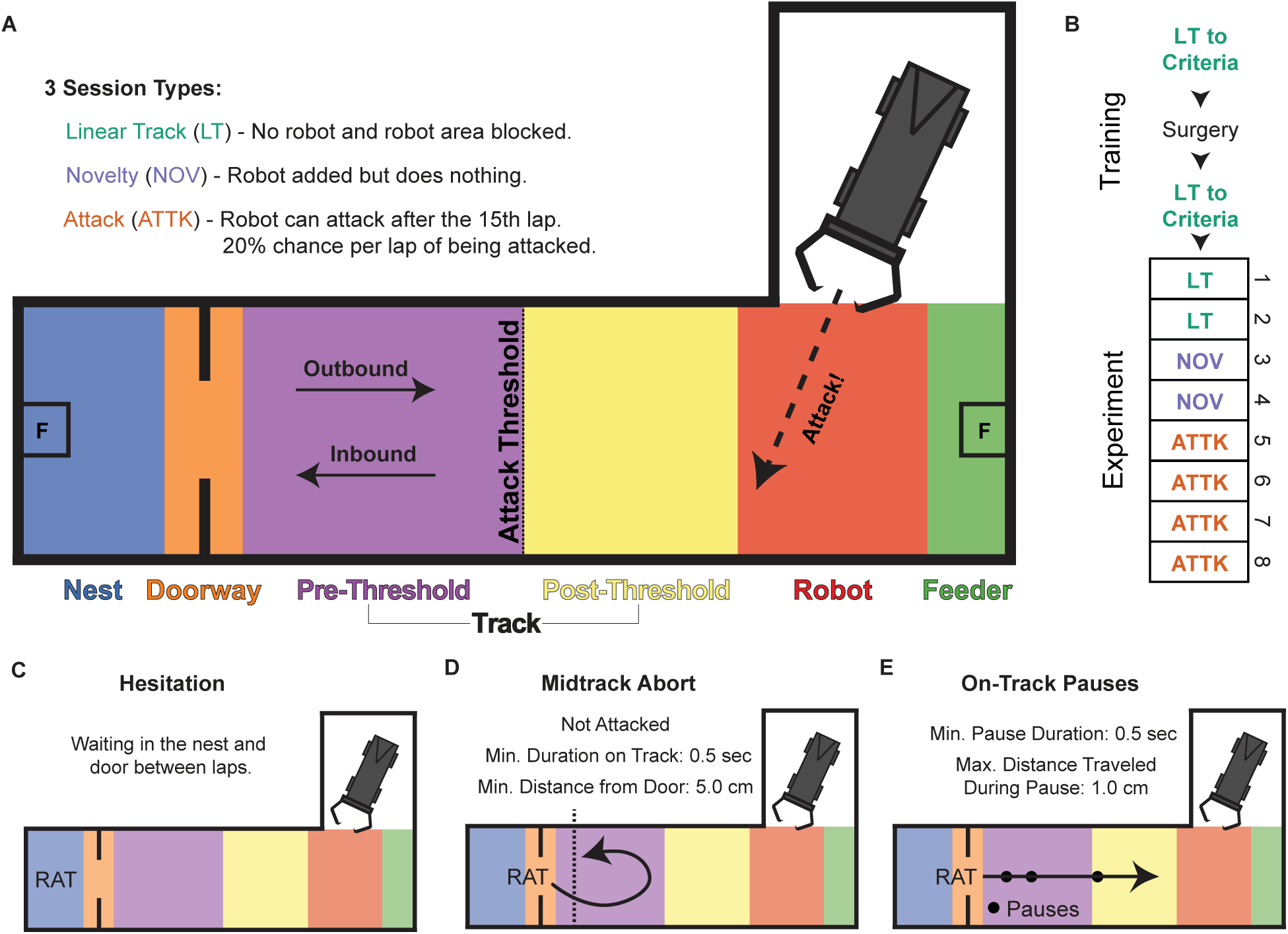
Task and experimental design. **A.** The track was composed of a nest, track, and robot bay in the shape of an L. There were two feeders (marked as F). During Novelty and Attack sessions a robot was placed in the robot bay, which was walled off during Linear Track sessions. During Attack sessions, there was a chance of the robot surging forth when the rat crossed the attack threshold. Based on these characteristics we divided the linear portion of the track into Nest, Doorway, Pre-Threshold Track, Post-Threshold Track, Robot, and Feeder segments. Journeys on the track were defined by the direction of travel when the rat entered the track; journeys that entered from the Doorway zone were Outbound and journeys that entered from the Robot zone were Inbound. **B.** The experiment consisted of 8 sessions: two linear track (LT), two novelty (NOV), and four attack (ATTK) sessions. Prior to the experiment, the rat was trained to alternate between feeders. **C, D, E.** Putative worry behaviors that we examined. **C.** Hesitation was defined as a period of waiting in the nest between laps. **D.** Midtrack aborts were defined as a spontaneous retreat while approaching the robot, that was not precipitated by the robot attacking. **E.** On-track pauses were instances when the rat hesitated on the track.

Our experiment consisted of three session types. On *Linear Track* sessions [2 days], the robot was not present and the robot bay was walled off to create a purely linear track between the nest and far feeders. On *Novelty* sessions [2 days], the robot was present but did not move or make noise. On *Attack* sessions [4 days], the robot would occasionally (1 in 5 chance of attacking on each lap after the first 15 laps) screech, surge forward, and move its pincer-like jaws and stinger-like tail when the rat crossed an unmarked attack threshold as they moved from the nest to the far feeder. To assess habituation and learning, the Attack sessions were divided into two *Early Attack* sessions [first 2 of 4 days] and two *Late Attack* sessions [last 2 of 4 days], giving us four conditions to compare of two-sessions each (Figure 1B). Sessions were analyzed in pairs for statistical power. For analytic purposes, we separated the track into spatial locations of the nest, doorway, pre-threshold track, post-threshold track, the robot itself, and the far feeder, and we separated the rats’ movement on the track into outbound and inbound journeys that indicated when the rat entered the track while moving towards the feeder and the nest, respectively (Figure 1A).

Once the rats were attacked by the robot, we observed three identifiable behaviors that have often been seen in similar approach-avoidance conflict tasks and are putatively indicative of worrying about the robot [37–39,44,69]: *hesitation in the nest*, *midtrack aborts*, and *on track pausing*. Hesitation in the nest entailed increased time spent in the nest space and thus reduced the number of robot approaches (Figure 1C). Hesitation likely indicates worry about being exposed to threat outside of the nest space. Midtrack aborts were instances when the rat left the nest and door area into the track zone, but spontaneously returned before reaching the feeder on the other side of the robot (Figure 1D). Midtrack aborts likely indicate a change of mind from approach to avoidance behavior. Retreats from attacks were excluded from the identification of midtrack aborts as those retreats were a reaction to the robot attack rather than reflective of a proactive concern. Lastly, we noticed that the rats began to pause in their outbound journeys (Figure 1E). This increase in on-track pauses was seen on outbound but not inbound journeys, and likely indicate concern about the danger from the robot.

### Hesitation in the nest

In approach-avoidance conflict, it is likely that a rat will be reluctant to approach the threat and may choose to avoid it entirely (Figure 1C). Rats ran fewer laps during the early Attack sessions (Figure 2A), and increased time in the nest (Figure 2B). Rats had a tendency to remain in the nest longer in the second half of the session than the first half in all sessions (presumably due to satiation); however, after being attacked, even early in a given session, rats hesitated in the nest longer than on non-attack sessions. To differentiate early vs late behavior on each session, we split attack sessions (sessions 5-8) before vs after the first attack on that session. For non-attack sessions (sessions 1-4), we split the session on a yoked lap that was a shuffled match of the first-attack lap from one of the attack sessions, which thus maintained the average and variation of attack laps from the attack sessions. Thus, we analyzed behavior for each session as before and after the first attack or yoked lap. Rats hesitated more before journeys towards the robot in the early Attack sessions after being attacked during that session compared to both the non-attack (Linear Track and Novelty) sessions and the late Attack sessions (Figure 2B).

**Figure 2.**
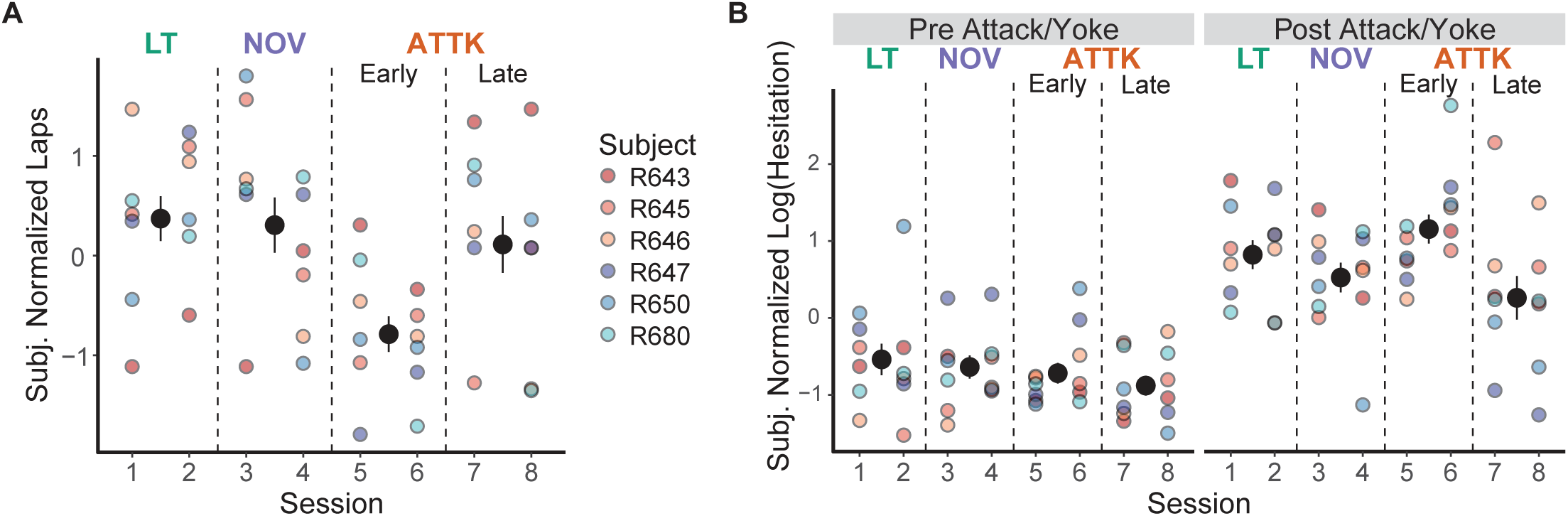
Effects of attack on hesitation behavior. **A.** Subject normalized laps run by the rat over the course of the experiment. Note that for all figures, the large dots with error bars are the mean and standard error of the subjects’ average behavior and the small dots are the individual subject values. **B.** The subject normalized log hesitation before and after being attacked. For sessions that the rat was not attacked, we yoked the pre-post split time to the beginning of the lap number that the rat was attacked during the attack sessions.

We statistically assessed this avoidance by fitting a linear mixed effects regression to the subject-normalized (in the statistical sense; see Methods) number of laps as a function of the session with a random effect of rat, and found a significant main effect of sessions, *F*(3, 39)=4.81, *p*=0.006. Rats ran significantly fewer laps on early Attack sessions than on Linear Track sessions, *t*(39)=-3.35, *p*<0.01, and Novelty sessions, *t*(39)=-3.16, *p*=0.02, but not on late Attack sessions, *t*(38)=-2.60, *p*=0.06. Rats did not run significantly fewer laps on late Attack sessions compared to Linear Track sessions, *t*(39)=-0.75, *p*=0.88, and Novelty sessions *t*(39)=-0.56, *p*=0.94. Rats ran similar numbers of laps on Novelty and Linear Track sessions, *t*(39)=0.19, *p*=1.00.

Rat hesitation in the nest showed a similar pattern. We found a significant interaction of session with pre/post attack/yoke, *F*(3, 2761)=6.46, p<0.001 and significant main effects of session, *F*(3, 2761)=6.66, p<0.001 and pre/post attack/yoke, *F*(1, 2762)=382.45, p<0.0001.

Post-hoc comparisons indicated that rats hesitated significantly longer during all sessions after the attack/yoke, all *t* scores > 8.5, *p*s<0.0001. During the pre attack/yoke period we did not detect significant differences in hesitation across the sessions, all |*t|* scores(2761) < 1.59, *p*s<0.38. However, during the post attack/yoke period there were significant differences across the sessions with more hesitation during the early attack sessions (vs. Linear Track: *t*(2761)=4.21, *p*<0.001, vs. Novelty: *t*(2761)=5.06, *p*<0.0001, vs. Late Attack: *t*(2761)=6.56, *p*<0.001). We found no significant differences in the duration of hesitation across Linear Track, Novelty, and late Attack sessions, all |*t|* scores <2.47, *p*s>0.06.

These results suggest that early attacks by the robot caused greater hesitation, but that the rats habituated to it in the later Attack sessions. Overall, the rats showed an increase in hesitation at the nest after first being attacked, which led to a reduction in the number of laps that they ran when they began to be attacked, but, in both cases, they habituated to the threat over the Attack sessions. In-nest hesitation may reflect the safety of the nest and active avoidance of the on-track threat.

#### Threat caused shift In spatial tuning of hippocampal cells

Place fields are thought to encode episodic memories [48,70] and have been observed to change in response to those events, encoding the location of the animal at the time of those events [71–73]. While previous work has found changes to spatial tuning in response to attack by the robot on similar tasks, in those previous task versions, the robot attacked towards the food pellet and the analyses could not differentiate the location the animal was at when attacked from the location of the robot itself or that of the reward site [46]. Previous work on the version of the task used here (linear track with a guarding robot attacking from the side) found a trend that spatial tuning was more stable on the nest side than the robot side [2].

The first attack by the robot produced a notable change in the spatial tuning of hippocampal place cells. These changes occurred in representations of space beyond the location of the animal when attacked, and instead appeared at the location of the robot and the approach to it (which was not where the animal was when attacked). Figures 3A and 3B show two examples of this shift in which a cell’s place fields adjusted to more heavily represent the space between the attack threshold and the robot. In the first example (Figure 3A), the cell’s spiking shifts from primarily covering the feeder to the space between the attack threshold and the robot. In the second example (Figure 3B), a cell with two place fields shifts to more heavily represent the approach to the robot.

**Figure 3.**
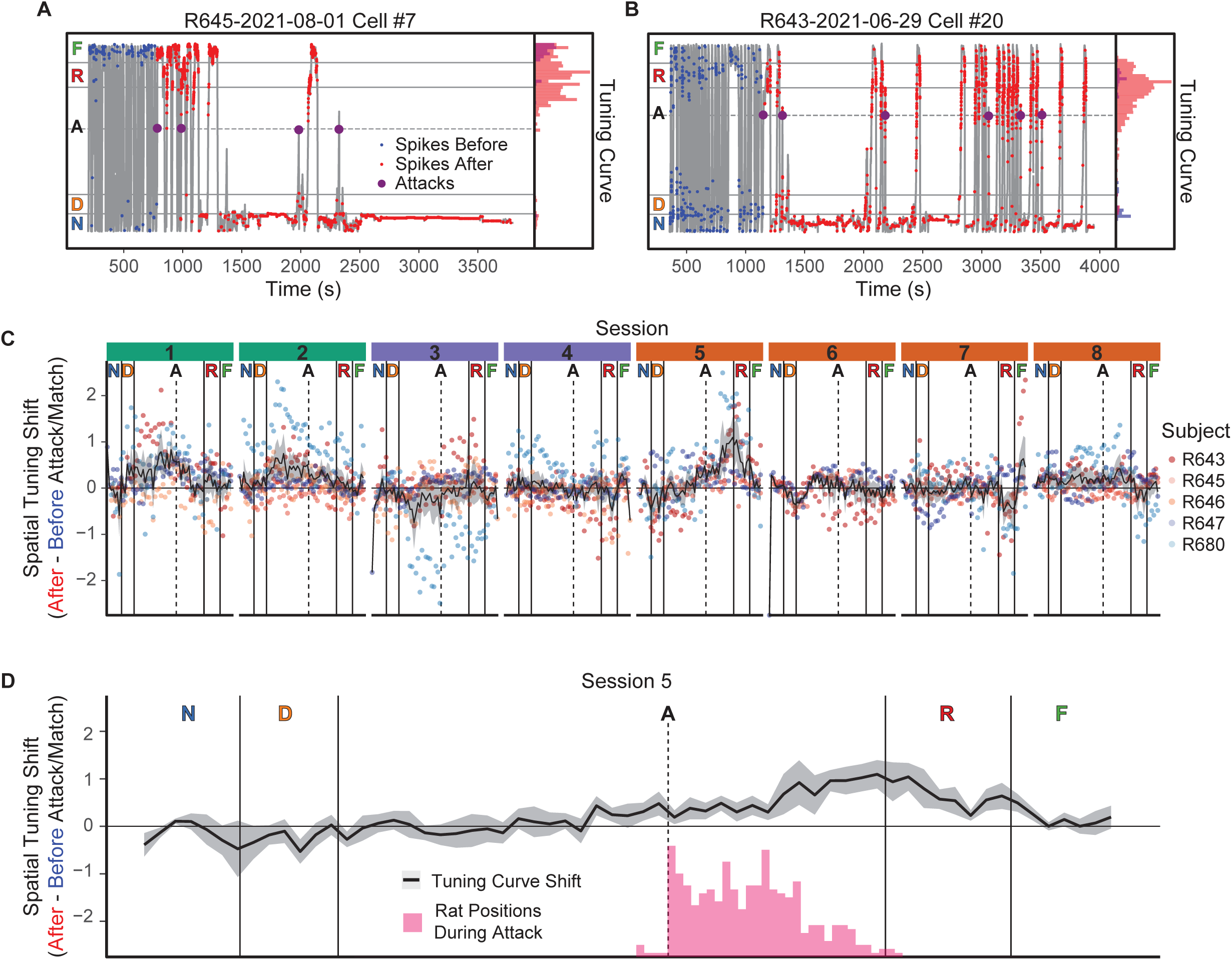
Effects of attack on spatial tuning. **A, B.** Example cells that developed place fields between the point of attack and the robot after being attacked the first time. **C.** The change in spatial tuning (i.e., the average change in cell tuning curves) before and after being attacked. The black line with gray fill indicates the average spatial tuning at each position and the standard error, and the small dots are each subjects’ shift at each tuning-curve spatial bin on the maze. Values above 0 indicate more spatial tuning to the location after the attack and values below 0 indicate less. Note the shift in representation on the first attack session (session 5) after being attacked around the location of the robot. **D.** The position of the rat 0 to 0.5 after the attack was initiated compared with the spatial tuning shift.

To quantify how spatial tuning adjusted to reflect the robot’s location, we determined the occupancy-normalized tuning curves based on each cell’s spiking activity prior to and after the first attack, and calculated the difference to get the occupancy-normalized spatial shift for each cell. After calculating this by cell, we averaged across the cells to establish the average change in spatial tuning at each location (Figure 3C). For sessions during which the rat was not attacked, we used the yoked laps as defined above. On the first attack day (session 5), there was a marked shift in the spatial tuning after the first attack in the robot and post-threshold track spatial segments.

We statistically assessed the shifts in spatial tuning by running a general linear model of the cell tuning curves as a function of the session and track zone. The GLM indicated that for the tuning curves on the approach to the robot there was a significant interaction of session and zone, *F*(35, 4189)=7.16, *p*<0.0001, a main effect of zone, *F*(5, 4189)=6.83, *p*<0.0001 and a main effect of session, *F*(7, 4189)=12.85, *p*<0.0001. Tukey post-hoc comparisons indicated that there was a greater shift of spatial tuning in the post-threshold track and robot zones on the first attack session (Session 5) than on other sessions(post-threshold track: |*t*| scores >5.42, *p*s<0.0001; robot: all |*t*| scores >3.27, *p*s<0.02). There was, also, significantly more representation of the pre-threshold track zone in the yoked post-attack duration during the linear track sessions than during other sessions, all |t|s>3.98, p<0.01. These results suggest that the attack by the robot caused the spatial tuning to shift to the robot and the approach to it.

Importantly, the change in the spatial representation was not directly reflective of the position of the rat during the attack (Figure 3D). While the positions of the rat during the 0 to 0.5 seconds after the attack is initiated are after the attack threshold, they do not reflect the largest change in spatial tuning, which is at the boundary of the track and robot sections of the linear track.

Overall, these results suggest that the rat’s hippocampus shifted its representation to the robot and the approach to the robot after it was attacked the first time. Importantly, unlike previous observations of place field appearances [71–73], these shifted representations encoded non-local locations (the robot and the final approach to it) rather than the location of the rat when the event occurred.

#### Threat representation during hesitation

As discussed in the introduction, theories suggest that worry entails representations of potential threats [1–4]. Given that dorsal hippocampal representations encode information about other places and other times [48,55,58,59,74,75], particularly in conditions of concern [36,56], we assessed the effect of the changes in hippocampal spatial tuning on the transient representations during HSEs during the increased hesitations after attack. If HSEs are used to replay information and there was a need to represent a salient event, then the rat’s hippocampus could represent the threat by increasing the number of sharp-wave ripples or by increasing the representation of the threat during the event.

During the time spent hesitating in the nest and at the nest entrance, CA1 LFPs had low theta power, instead showing increased delta (1-4 Hz) and SWR (150-250 Hz) power, indicative of large-amplitude irregular activity (Figure 4A; LIA, [47–49]). SWRs were also indicated by cross-frequency correlations, which highlight a block of highly correlated frequency power in the SWR band (Figure 4B). We decoded during HSEs, which overlaps with most SWRs (Figure 4C). We did not see a marked increase in the rate of SWRs (Figure 4D) or HSEs (Figure 4E; *F*(3,11)=2.49, p=0.11) while the rats were in the nest after that first attack. We did detect a main effect of session with the SWR rate, *F*(3,12)=3.66, p=0.04, but none of the post-hoc comparisons survived correction for multiple comparisons, all |*t*|-scores < 2.88, *p*s>0.05.

**Figure 4.**
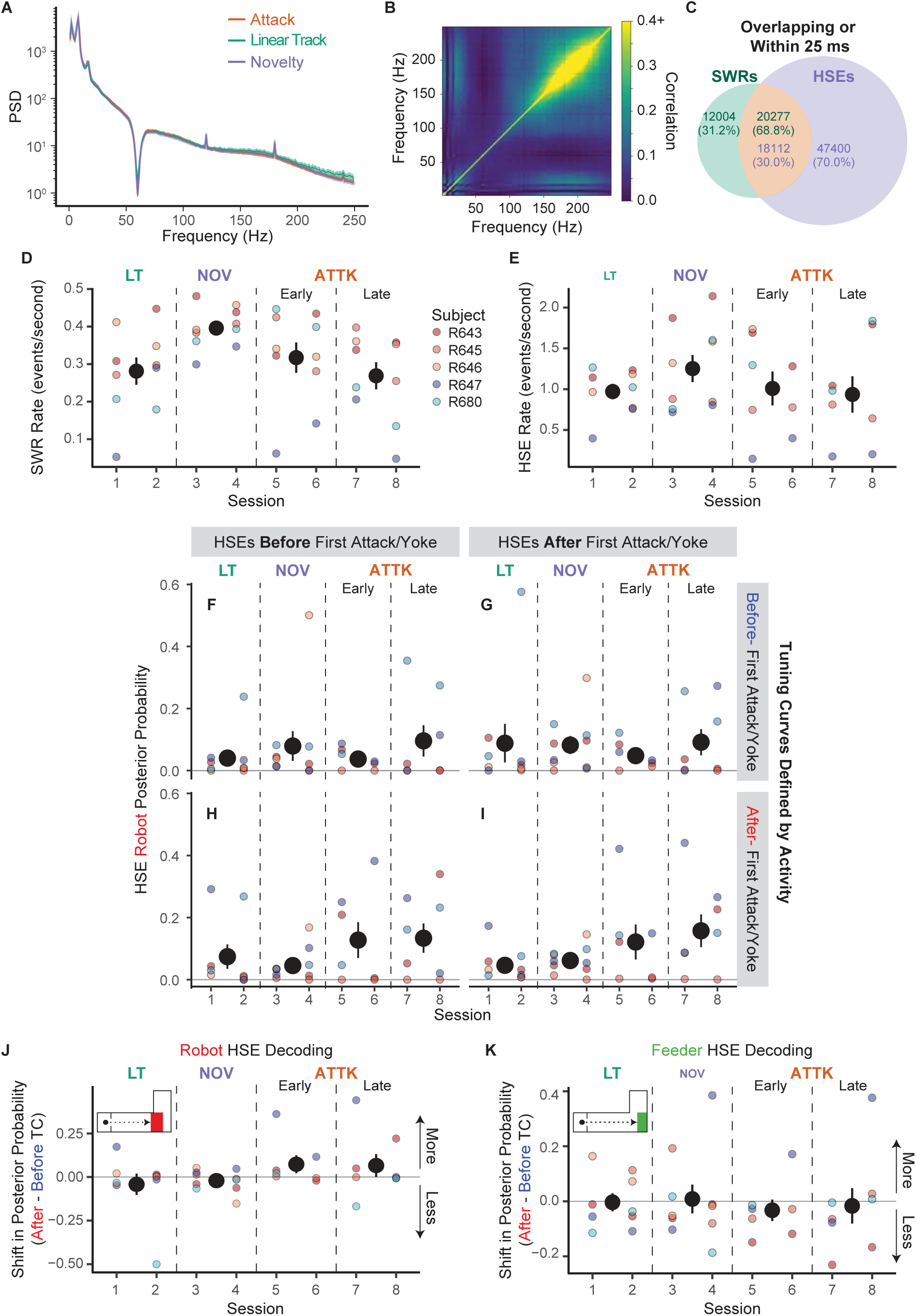
Changes in representation during sharp-wave ripples (SWRs) and high-synchrony events (HSEs). **A.** The average power spectral densities while rats were in the nest across sessions. **B.** The average cross-frequency correlation, which indicates significant sharp-wave ripples (150-250 Hz). **C.** The overlaps in SWRs and HSEs. The numbers indicate the event counts and, in parentheses, the percentiles. The different colors indicate whether the overlap is determined based on the SWRs (green) or HSEs (purple). **D.** The SWR rate while the rats were in the nest. **E.** The HSE rate while the rats were in the nest. **F-I.** The posterior probability of the robot being represented. The panels are separated by whether the HSEs were before or after the first attack/yoked time and whether the decoding was performed based on tuning curves from before or after the first attack/yoke. These panels suggest that the shift in cell tuning curves mediates the increased robot representation while rats hesitated in the nest. **J.** The difference in robot posterior probabilities between applying before- and after-attack tuning curves to decode hippocampal representation. **K.** The difference in feeder posterior probabilities between applying before- and after-attack tuning curves to decode hippocampal representation.

Importantly, the content encoded in the neural ensembles also could have changed. Earlier work indicated an increase in representation of the robot location during SWR events during hesitation in the nest after the rat had been attacked [2]. We found a similar increase in representation of the robot location during high synchrony events during hesitation after the rat had been attacked, χ^2^(1)=260.07, *p*<0.0001.

Further analysis, however, found that this was due to the change in spatial tuning — not to a change in the cells activating during the HSE. An increased representation of the robot could arise either from a change in which cells participate in the HSE (due, for example, to what aspects of the world are attended to during those HSEs), or by a change in the underlying representation of the world (due, for example, to a change in the place fields). Our data definitively show the latter.

To determine what was causing the increased representation of the robot’s location during HSEs, we compared four conditions: decoding of HSEs occurring before and after the first attack (or yoked lap) crossed against spatial tuning derived from experiences occurring before and after the first attack (or yoked lap) (Figure 4F-I). We observed that there was more representation of the robot when we used the tuning curves that were defined by activity after being attacked (Figure 4H and Figure 4I), but not when we used the before-attack tuning curves (Figure 4F and Figure 4G), even for HSEs that occurred before the rat was attacked (Figure 4H). This suggests that robot representation is mediated by shifts in spatial tuning.

We statistically assessed this shift by fitting a general linear mixed effects regression to the posterior probability of the robot during after-attack (or post matched yoked lap) high synchrony events as a function of whether the rat had been attacked that session or after the matched (yoked) lap and the session of the experiment with a random factor of rat. The general linear mixed effects regression detected a significant interaction, χ^2^(3)=946.43, p<0.0001 and main effect of session, χ^2^(3)=365.40, p<0.0001, but no main effect of whether we used the pre-attack/matched lap tuning curves, χ^2^(1)=0.00, p=0.997. Post-hoc comparisons indicated that when using the after-attack tuning curves, there was more decoding of the robot during the early and late Attack sessions than the Linear Track and Novelty sessions (early Attack to Linear Track: z=15.94, p<0.0001, early Attack to Novelty: z=7.16, p<0.0001, late Attack to Linear Track: z=27.17, p<0.0001, late Attack to Novelty: z=22.45, p<0.0001). There was also significantly less decoding of the robot on Linear Track than on Novelty sessions, *z*=-11.76, p<0.0001. When using the pre-attack/lap-matched tuning curves, we no longer detected this relationship. With pre-tuning curves, there was no detectable difference between the decoding during early Attack and the Linear Track sessions, z=1.96, p=0.20 and less representation than during Novelty sessions, z=-11.11, p<0.0001. Similarly there was less decoding of the robot during late Attack sessions than during the Linear Track, z=-4.45, p=0.0001, and Novelty sessions, z=-17.31, p<0.0001.

For comparison, we assessed representation of the feeder during these same high synchrony events and found the opposite pattern (differences summarized in Figure 4K). When using the post-attack tuning curves, there was significantly less representation of the feeder during early and late Attack sessions than on Linear Track and Novelty sessions (early Attack vs Linear Track: z=-7.79, p<0.0001, early Attack vs Novelty: z=-2.995, p=0.01, late Attack vs Linear Track: z=-8.519, p<0.0001, late Attack to Novelty: z=-3.304, p<0.01). When using the pre-attack tuning curves, there was more representation of the Feeder during early and late Attack sessions than on Linear Track and Novelty Sessions (early Attack vs Linear Track: z=15.99, p<0.0001, early Attack vs Novelty: z=11.33, p<0.0001, late Attack vs Linear Track: z=19.89, p<0.0001, late Attack vs Novelty: z=15.57, p<0.0001).

These results indicate that there was increased representation of the robot during high-synchrony events occurring while the rat was in LIA, hesitating in the nest, after the animal was attacked, but that this increased representation did not arise from an increased probability of events being released, nor from a change in the set of cells firing during the event. Instead this increased representation came from changes in place field encoding. The hippocampal spatial tuning had shifted to more highly represent the location of the robot and thus HSEs were more likely to pull up that representation. The hippocampal map of the world had changed to emphasize the location of the robot, and thus the mental explorations of the world during HSEs were now changed to include the robot.

### Midtrack Aborts

During an approach-avoidance conflict, an individual may initially decide to approach a reward, but then be overwhelmed by the potential threat as it becomes more salient and switch to an avoidance strategy. On this task, this change of strategy manifests as a midtrack abort [37–39,44] (Figure 1D). Consistent with this previous work, after being attacked, rats began to engage in more midtrack aborts when approaching the robot (Figure 5A). We did not see a similar change in the number of reorientations on journeys back to the nest (i.e., away from the robot; Figure 5A), implying that the increase in these midtrack aborts were selective to the approach-avoidance conflict that occurs on the outbound journey towards the robot.

**Figure 5.**
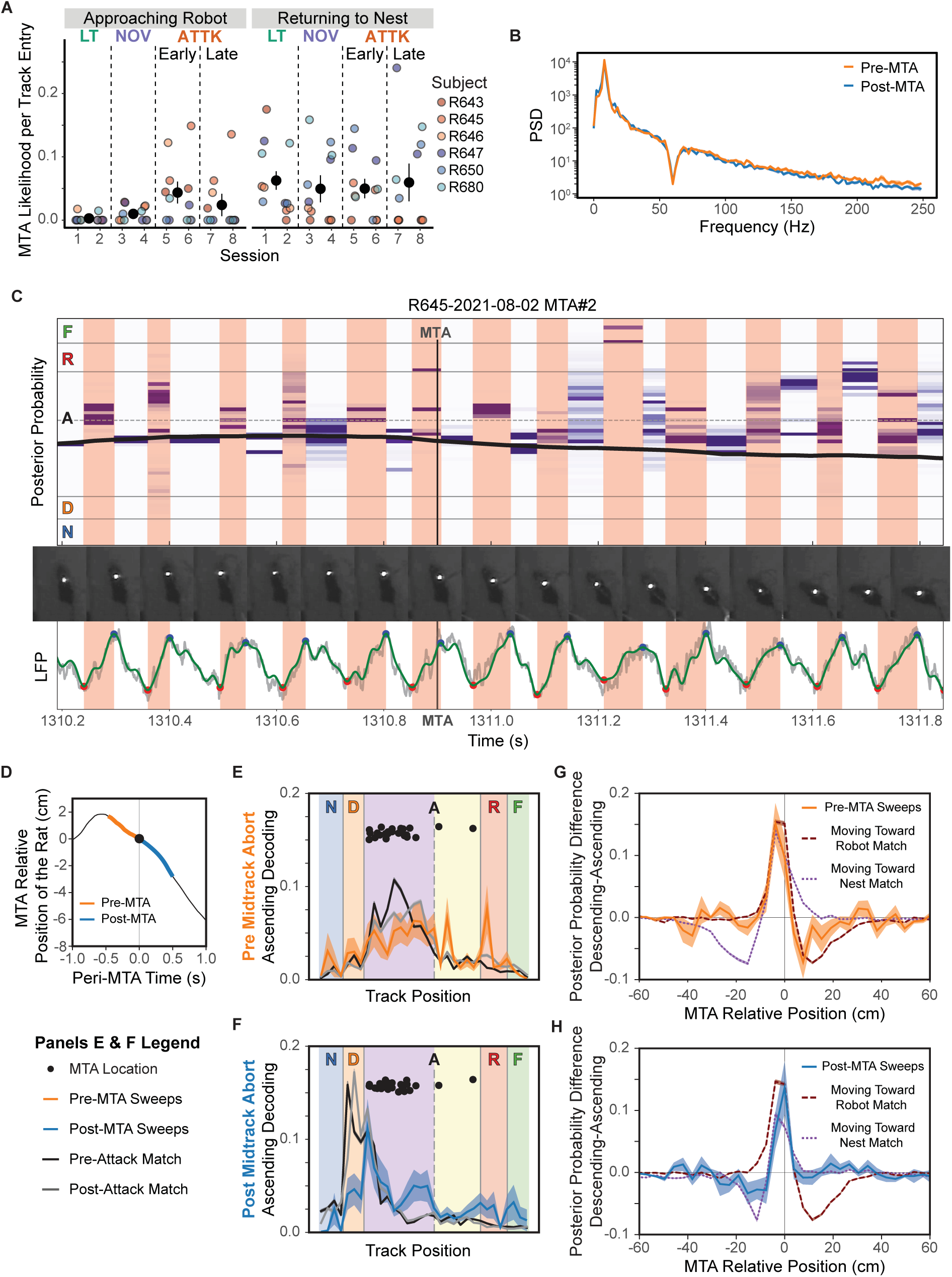
Midtrack aborts (MTA). **A.** Probability of a midtrack abort per entry onto the track. **B.** the power spectral density before and after a midtrack abort. **C.** Example of how the decoded posterior probability changes around the midtrack abort, animal’s position indicated by (the black line. The frames below the decoding highlight the rats movement around the MTA. The LFP shows the identified theta peaks (blue) and troughs (red), which provide the ascending (red background) and descending (white background) phases of theta. The green line shows 6-40 Hz bandpass filtered local field potential. **D.** The average position of the rats around a midtrack abort (black), with the pre-attack and post-attack periods of investigation respectively highlighted orange and blue. **E.** The posterior probability during the ascending phase of theta before the midtrack abort. **F.** The posterior probability during the ascending phase of theta after the midtrack abort. **G.** The direction of the decodings from the descending to ascending phases of theta before the midtrack abort. **H.** The direction of the decodings from the descending to ascending phases of theta after the midtrack abort was initiated.

We statistically assessed this spontaneous switch from approach to avoidance behavior by fitting a binomial regression to whether the animal aborted during a journey as a function of session and direction of travel. This analysis found that there was an interaction of direction of travel and session, χ^2^(3)=21.87, p<0.001, a main effect of session, χ^2^(3)=16.87, p<0.001, and a main effect of direction, χ^2^(1)=21.87, p<0.001. When approaching the robot, the rats made significantly more midtrack aborts on early Attack sessions than during Linear Track sessions, z=3.75, p=0.001, and Novelty sessions, z=3.52, p=0.002. This was somewhat attenuated during the late Attack sessions, which were not significantly different from early Attack sessions, z=1.94, p=0.21, and more likely to spontaneously abort an approach than during Linear Track sessions, z=2.75, p=0.03, but did not significantly differ from Novelty sessions, z=1.80, p=0.27. As expected, the rats did not significantly differ in the number of midtrack aborts during Linear Track and Novelty sessions, z=-1.54, p=0.41. There were no significant differences in the number of midtracks aborts across the sessions when returning to the nest (all |z| scores <= 1.5, ps>0.43).

Overall, the rats showed an increase in the number of midtrack aborts on outbound journeys. This increase was selective to the approach-avoidance conflict as indicated by the increase only occurring when approaching the robot on outbound journeys.

#### CA1 representations swept to the attack threshold during midtrack aborts

Extensive previous data has found that as rats are running towards a goal, the dorsal hippocampus shows theta oscillations [47,52]. The first half of the theta cycle encodes the location of the animal and the second half encodes potential future paths, sweeping forward towards the next goal [9,24–28,53,76]. If hippocampal information of a threat’s location is being used to determine whether the animal should retreat as it approaches, then that information will most likely be observed during the second half of theta oscillations. Figure 5C shows an example of how the forward sweep decoding changed around the time of a midtrack abort. In this example, the decoding posterior probability during the descending component of theta (shaded white) encoded the location of the animal quite accurately (purple line). In contrast, during the ascending component of theta sweeps (shaded red), the decoding posterior probability was concentrated substantially ahead of the animal’s position prior to the MTA. This you-are-here / sweep-forward structure is consistent with theta sequences seen in goal-approach (non-threat-based) tasks [25,27,53,77–80].

However, after initiating the midtrack abort, these local and forward representations during theta became more varied. There were no apparent changes in the low-frequency power spectral density before vs after the midtrack abort (Figure 5B; no significant differences in 1-4 Hz delta, *t*(41)=-2.37, p=0.02, 6-10 Hz theta, *t*(41)=0.57, p=0.57, or 15-20 Hz beta, *t*(41)=-1.43, p=0.16, after a family-wise error correction), although there was a significant decrease in 120-250 Hz sharp-wave ripple power band power, *t*(41)=3.03, p=0.004. Given the observed power spectral density, this SWR-range significant difference may have been due to changes in the aperiodic component rather than oscillatory activity [81], as no sharp-wave ripple events were observed. These analyses suggest that the hippocampus remained in the theta state during midtrack aborts.

To determine the relationship between the hippocampal representations and the behavior around the midtrack abort, we decoded the information represented during ascending theta phase 0.5 seconds before and after the rat the MTA event (Figure 5D). We aligned the data to the moment when the rat’s head began to turn into the retreat, which typically occurred after the rat had already stopped their forward momentum. We compared the forward theta component around the MTA to when the animal was in the same position and moving in the same direction. We separated our comparisons by whether the animal had been attacked yet in a given session since that affected the rats’ movement rates and, likely, their action planning.

The ascending theta decodings around the MTAs were atypical to matched comparisons (Figure 5E and 5F). Prior to the MTA, we observed more decoding of the robot and the attack threshold than matched comparisons (Figure 5E). The representations prior to the MTA tended to be more varied than the matched comparisons. Importantly, the ascending theta decodings we observed after the MTA were unlike their matched counterparts (Figure 5F). The second half theta cycle decodings in the 0.5 s after the MTA became more like when moving towards the nest, but still included greater representations of the robot and the attack threshold, both of which were behind the rat (Figure 5F).

Since a mid-track abort (MTA) is a switch in the rats’ direction of travel, the local-to-ahead direction of the theta decoding should track the anticipated action. We assessed this by comparing the five theta sweeps before and after the MTA with position and direction matched controls. By subtracting the decoding during the ‘local’ descending phase of theta from the ‘forward’ ascending phase we could assess how much of the theta representation was local and how much was non-local (Figure 5G). We assessed this by decoding the descending (you-are-here) portion (shaded white in Figure 5C) and ascending (sweep-forward) portion (shaded red in Figure 5C) separately and subtracting them (descending - ascending). This typically shows a clear distinction between the two representations [27,82]. Despite the rat being stationary, the second half of theta had significant forward representation towards the outbound journey (Figure 5G). Unlike sweeps on approach-approach conflict tasks during vicarious trial and error events, in which the sweeps go farther ahead than during matched passes that don’t include vicarious trial and error events [9,24,25], these sweeps were more attenuated than matched passes. This could be due to the sweeps only going to the attack threshold or the robot location rather than all the way to the far feeder (Figure 5E). Similarly, once the rat had started their retreat, the direction of the theta decoding tracked the rat’s future direction of travel back to the nest (Figure 5H), even though the rat had not fully turned to face that direction (see, for example, Figure 5C). Importantly, however, the ascending (sweep “forward” to the nest) components were attenuated forward (toward the nest) and included components behind the rat (where the robot was; Figure 5H), suggesting an attention to dangers even behind the rat during these theta states.

### On-track pauses

We observed that the rats began to repeatedly pause as they moved towards the robot after it had attacked them (Figure 1D, compare Figure 6A with Figure 6B). We observed a marked increase in the number of pauses on outbound journeys when the rat was approaching the robot after being attacked (Figure 6B). During non-attack sessions the rats rarely paused when approaching the feeder, but this on-track pausing behavior increased dramatically after the rat was attacked. There was a decrease of on-track pausing on inbound journeys, suggesting that these pauses were related to the approaching danger of the robot attack.

**Figure 6.**
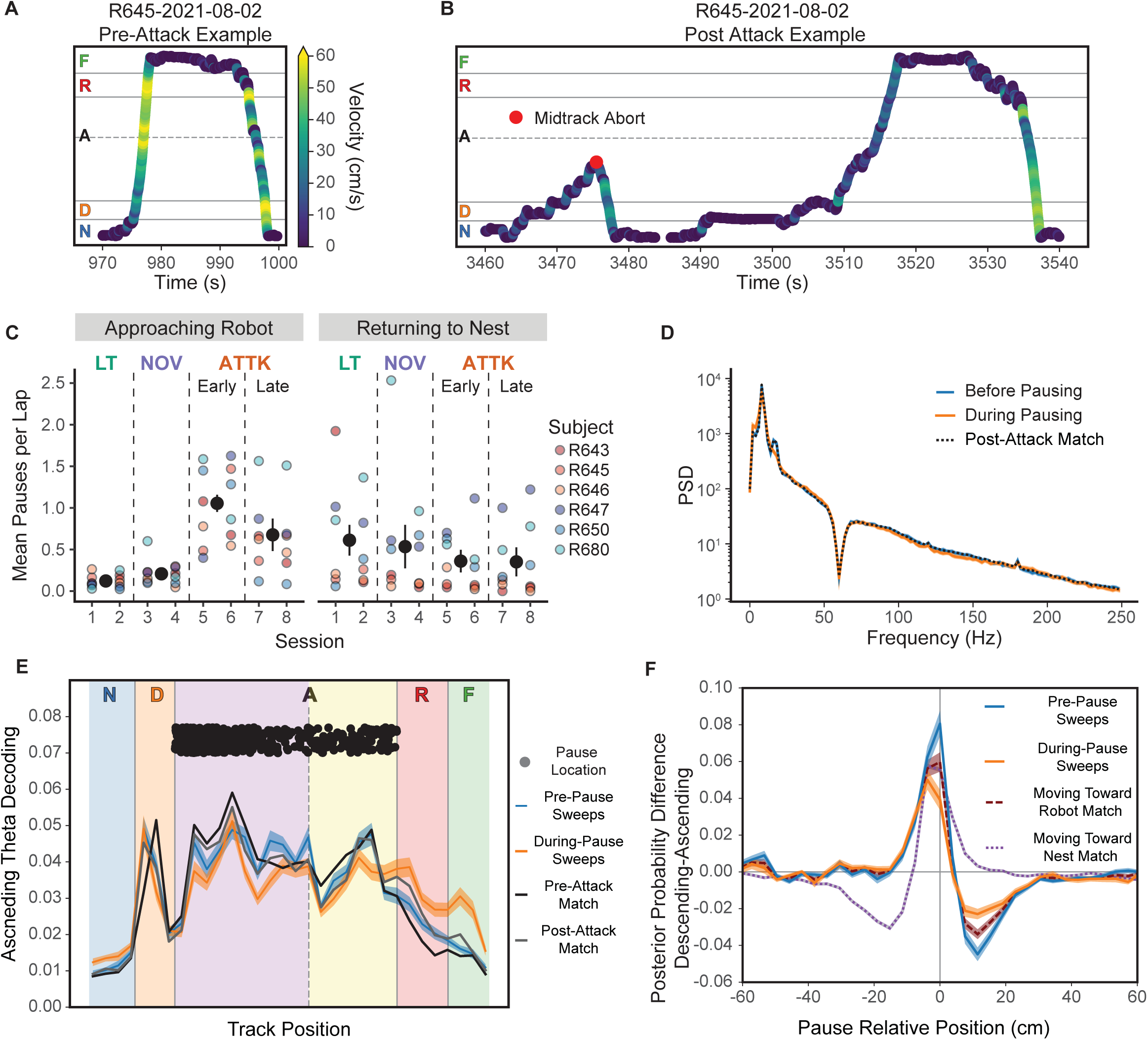
On-Track Pausing. **A.** Example rapid movement on a lap prior to being attacked (the first attack was at 1091 s). **B.** Example movement of the same rat that now shows more frequently pausing on the track after being attacked. **C.** The average number of pauses per lap during the approach to the robot. **D.** The power spectral density prior to and after the pause was initiated. **E.** Decoded posterior position during the ascending phase of theta (non-local) prior to pausing on the track. **F.** The direction of the decodings from the descending to ascending phases of theta prior to pausing.

We quantified these changes by fitting a generalized linear mixed effects regression to the number of pauses on a journey towards the feeder as a function of the session with a random factor of rat. We found a significant main effect of session, χ^2^(3)=631.60, *p*<0.0001.

Rats paused significantly more often on the outbound journey during the early and late Attack sessions compared to the Linear Track sessions (vs early Attack: *z*=19.43, *p*<0.0001; vs late Attack: *z*=14.79, *p*<0.0001) and Novelty sessions (vs early Attack: z=18.60, p<0.0001; vs late Attack: *z*=12.53, *p*<0.0001). We found increased pausing on the early Attack sessions compared to the late Attack sessions, *z*=8.33, *p*<0.0001, consistent with habituation. Rats also paused less on Novelty sessions than on Linear Track sessions, *z*=-4.37, *p*=0.0001, suggesting that the pauses were due to being attacked, and not simply to the presence of the robot.

The return journey to the nest provides an informative comparison for the effect of the threat (Figure 1A) as it should have an opposite effect. The rats should become more eager to return to the safety of the nest after being attacked. We observed that it did (Figure 5C). As predicted, rats showed a decrease in pausing on the return after being attacked. When we statistically assessed this relationship we found a significant main effect of session, χ^2^(3)=69.13, *p*<0.0001. Rats paused significantly less on early and late Attack sessions than on Linear Track (vs early Attack: *z*=-5.55, *p*<0.0001; vs late Attack: *z*=-5.66, *p*<0.0001) and Novelty sessions (vs early Attack: *z*=-6.04, *p*<0.0001; vs late Attack: *z*=-6.18, *p*<0.0001). There was not a significant difference in the pausing rate between early and late Attack sessions, *z*=-0.17, *p*=1.00, nor between Linear Track and Novelty sessions, *z*=-0.49, *p*=0.96.

Rats began to show these on-track pauses only after being attacked by the robot and not simply by its presence — on-track pausing did not increase during the Novelty sessions, but only during the early Attack sessions. Rats continued to show on-track pausing during all four Attack sessions, but did show some habituation between the early and late Attack sessions.

Importantly, not only did rats not show on-track pauses on the inbound journey back to the nest, they actually showed a decrease in pausing during that inbound journey, suggesting that they may have been running faster to return to the safety of the nest.

#### CA1 remained in theta during on-track pausing

If the on-track pauses are considerations of potential future danger (like MTAs), we would expect CA1 to remain in theta and show representations of the robot (Figure 5B). If the on-track pauses are moments of hesitation, we would expect CA1 to be in LIA and show SWRs and HSEs that represent the robot (Figure 4A). While we did find a decrease in 6-10 Hz theta power (*t*(609)=-7.12, p<0.0001), an increase in delta power (*t*(609)=5.48, p<0.0001), and a major decrease in 15-20 Hz beta power (*t*(609)=-17.55, p<0.0001), theta power remained relatively strong throughout the pause (compare Figure 6D to Figure 4A). We also saw a significant decrease in SWR power during these pauses (t(609)=-5.62, p<0.0001) and a significant decrease in HSE count (*t*(14)=3.49, p=0.004), strongly suggesting that CA1 remained in relatively normal theta through the pause.

We also found some evidence for increased CA1 representation of potential threats and rewards during on-track pauses (Figure 6E). During pauses the ascending theta sweeps represented the robot and feeder more than matched comparisons or the pre-pause sweeps. To determine whether anything in the hippocampal representations might have initiated the on-track pause, we assessed the representation during the decoded ascending (“sweep-forward”) component of theta (Figure 6F). There tended to be a larger difference between the descending and ascending phases prior to pause than during the pause. These data suggest that pauses are somewhat similar to MTAs in that there is greater representation of the robot. Importantly, the increase in feeder representation during the pause (Figure 6E) as compared to the pre-MTA feeder representation (Figure 5E) may be what differentiates the two actions (continuing forward vs. retreating).

### Pharmacological manipulations with diazepam

Diazepam is a well-studied and frequently prescribed anxiolytic. Previous work has found that diazepam mediates behavioral measures of anxiety on this task, including changes in hesitation time and number of midtrack aborts [39]. With a separate cohort of rats (n=5, 3 M, 2 F), we administered diazepam or vehicle (Tween-20, see methods) intraperitoneally (IP) to rats on alternating Attack sessions (Figure 7A). These rats were first trained to find food at alternate locations (nest and far feeder) as with the previous experiment, and then were given two days of saline injections to get them used to receiving IP injections and while they continued to run the linear track each day. The robot was present, but not attacking on these two saline days, making them Novelty sessions. The rats then went straight to the Attack sessions, alternating diazepam or vehicle, counterbalanced across rats. Single shank silicon probes were implanted unilaterally into CA1 before the first Saline session. Unfortunately, these probes had higher impedances and, thus, did not permit single cell identification but did allow us to assess the local field potential.

**Figure 7.**
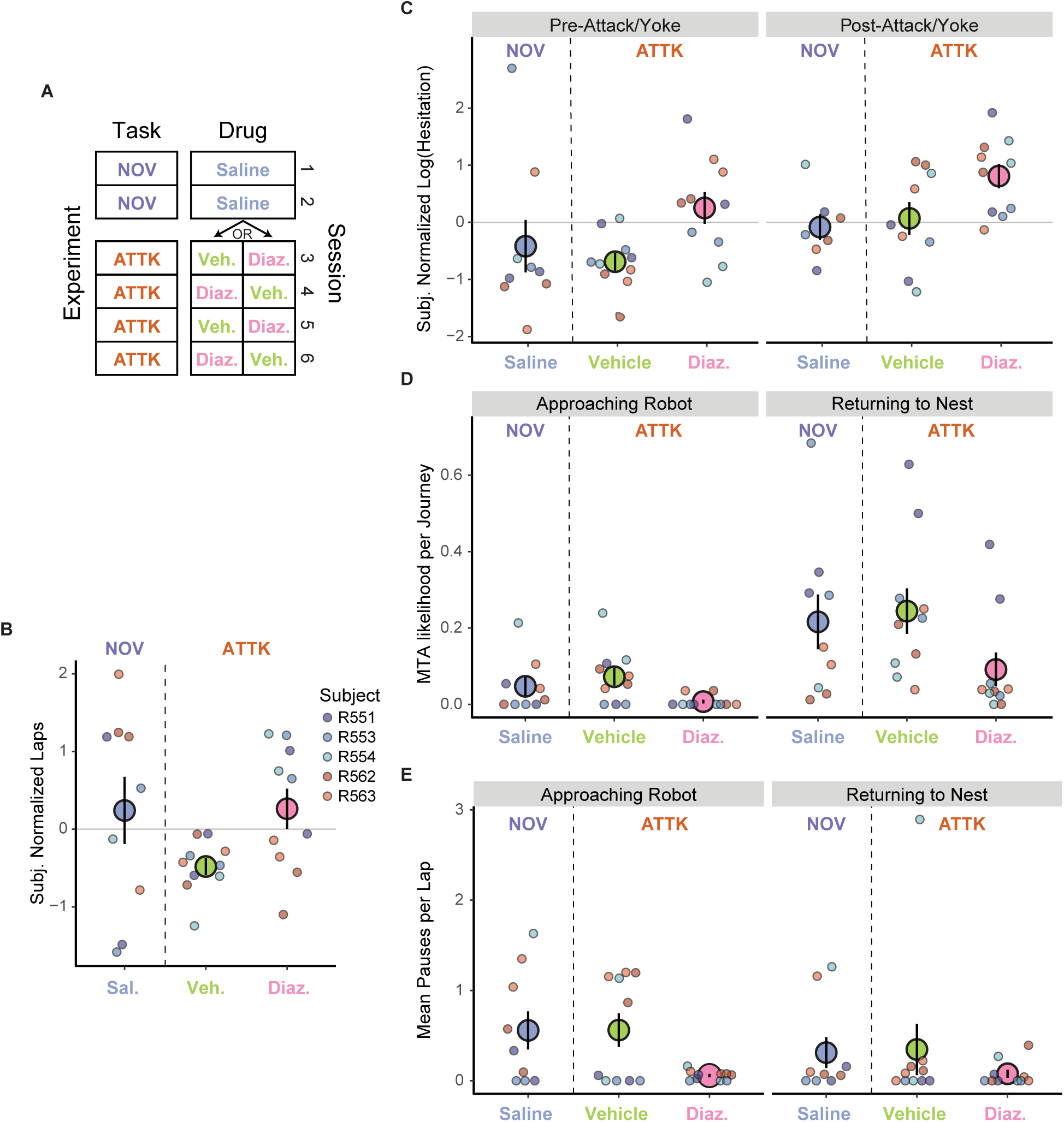
The effects of diazepam on behavior. **A.** This experiment consisted of 6 sessions: two linear track (LT), on which the animal received saline injections, and four attack (ATTK) sessions, on which the animal received alternating diazepam or vehicle (Tween20) injections, with half of the rats starting with vehicle and the other half with diazepam. **B.** The effect of diazepam and threat on the number of laps that the rats ran. **C.** The effect of diazepam and threat on hesitation before and after the rat was attacked. **D.** The effect of diazepam and threat on the probability of the rats engaging in a midtrack abort. **E.** The effect of diazepam and threat on the number of pauses while approaching the robot.

Diazepam had similar behavioral effects to that seen in previous work [39], and were consistent with the expected effects of an anxiolytic. Specifically, we observed that when administered diazepam on Attack sessions, the rats ran more laps than the Tween-20 controls (Figure 7B; *t*(14)=2.75, *p*=-0.02), hesitated longer in the nest (Figure 7C), were less likely to engage in a midtrack aborts (Figure 7D), and paused less frequently on the track (Figure 7E).

When rats were administered diazepam they hesitated longer and were not as strongly affected by threat as when they were only administered the vehicle (Figure 7C). We detected a significant interaction of administered drug with whether the rat had been attacked that session on hesitation, *F*(1, 342)=4.69, *p*=0.03, and main effects of the drug, *F*(1, 645)=40.13, *p*<0.0001, and whether the rat had been attacked, *F*(1, 453)=24.10, *p*<0.0001. When administered diazepam, the rats hesitated longer than when only administered the vehicle regardless of whether it was before, *t*(732)=6.32, p<0.0001, or after the first attack, *t*(267)=2.73, p<0.01. As a reaction to the first attack, when the rats were administered the vehicle they significantly increased their hesitation, *t*(571)=4.91, p<0.0001, but when administered diazepam they only trended to hesitating longer, *t*(252)=1.96, p=0.051.

Once the rats entered the track, however, they were less likely to perform a midtrack abort under diazepam (Figure 7D). We detected a significant interaction of administered drug with the direction of travel on midtrack aborts, χ^2^(1)=4.19, p=0.04, and main effects of the drug, χ^2^(1)=30.22, p<0.0001, and direction of travel on the track, χ^2^(1)=28.10, p<0.0001. When administered diazepam, the rats were less likely to perform a midtrack abort than when only administered the vehicle, regardless of the direction of travel (outbound: z=-3.92, p=0.0001; inbound: z=-6.44, p<0.0001). As with the first cohort (Figure 5A), rats were also more likely to engage in a midtrack abort when returning to the nest than when moving towards the robot (diazepam: z=3.86, p<0.0001; vehicle: z=-5.34, p<0.0001).

Similar to how diazepam affected midtrack aborts, it also reduced the number of pauses on the track (Figure 7E). When moving towards the robot and administered diazepam, the rats paused significantly less frequently than when only administered the vehicle, z=-9.35, p<0.0001. This was also the case when the rats returned to the nest, z=-10.08, p<0.0001.

#### Diazepam eliminated SWRs and changed theta asymmetry

In addition to its profound behavioral effects on anxiety-like behaviors (hesitation in the nest, midtrack aborts, and on-track pausing), diazepam had profound effects on the hippocampal processing in both theta and LIA states.

As can be seen in Figure 8A, the oscillatory increase in SWR power normally seen in hippocampal power spectral densities is absent under diazepam. During hesitation in the nest (Figure 8B), SWRs vanished (Diazepam vs. Tween: *t*(18)=-3.26, p=0.01). Because SWRs are transient events, small numbers of these events can be missed in power spectral densities; however, such events appear as a strong block of correlation in cross-frequency correlation plots [83]. Typically, SWR activity can be observed as cross-frequency correlations in the 150-250 Hz range, which can be seen when the rats were administered saline or vehicle. In contrast, when the rats were administered diazepam, these cross-frequency correlations vanished (Figure 8C).

**Figure 8.**
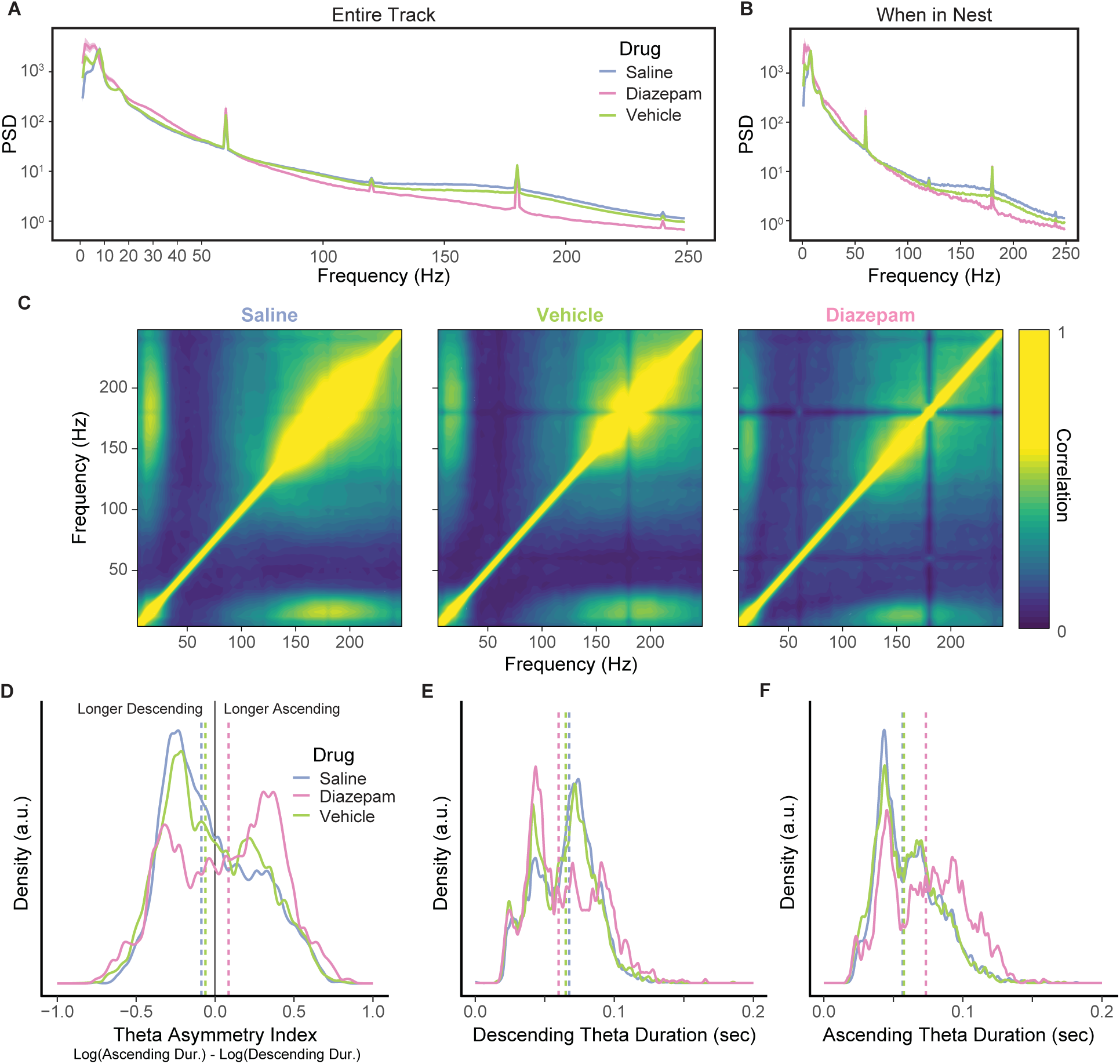
The effects of diazepam on local field potentials. **A.** The effect of diazepam on the average power spectral densities during the course of the sessions. **B.** The PSD when only looking at while the rat is in the nest. **C.** The reduction in the cross-frequency correlation of frequencies; note the missing correlation block indicated a lack of sharp-wave ripple events (150-250 Hz) when the rats were administered diazepam. **D.** The change in theta asymmetry with the administration of diazepam. The vertical dashed lines indicate the median value. **E.** How diazepam affects the duration of the descending phase of theta. **F.** How diazepam affects the duration of the ascending phase of theta.

LFPs under diazepam (Figure 8A) showed strong theta and beta power with an increase in 30-50 Hz gamma power, which likely implies changes in the balance between entorhinal and CA3 inputs [84,85]. Importantly, the peak frequency of theta, shifted discernibly from 8 to 6 Hz.

Diazepam significantly altered theta asymmetry during the outbound journeys (Figure 8D) by making the ascending period of theta longer. There was a significant main effect of the drug manipulation on theta asymmetry, χ^2^(2)=101.5, p<0.0001, and when the rats were administered diazepam those rats had significantly greater theta asymmetry (diazepam vs. saline: z=9.84, p<0.0001; diazepam vs. vehicle: z=8.59, p<0.0001). There was no difference in theta asymmetry between when the rats were administered diazepam and saline, z=1.14, p=0.49. The shift in theta asymmetry was primarily due to the duration of the ascending phase of theta increasing (non-local, Figure 8F), rather than the descending phase (local, Figure 8E). There was a significant main effect of the administered drug on the duration of the ascending phase of theta, χ^2^(2)=490.98, p<0.0001. When the rats were administered diazepam the duration of the ascending phase was significantly longer (diazepam vs. saline: z=20.59, p<0.0001; diazepam vs. tween: z=20.55, p<0.0001), and there was no difference when they were administered saline and vehicle, z=1.08, p=0.53. During the descending phase of theta (Figure 8E) there was, also, a significant main effect of the administered drug on the duration, χ^2^(2)=19.37, p<0.0001. While the descending phase of theta was longer in rats that were administered diazepam when compared to when administered the vehicle, z=4.09, p=0.0001, we did not detect a difference from when the rats were administered saline, z=1.85, p=0.15. Additionally, the length of the descending phase was longer when the rats were administered saline, than when administered the vehicle, z=3.18, p=0.004. This pattern of effects suggests that the threat of the attack makes the descending phase of theta longer, but that diazepam prevents this change.

## Discussion

Rats facing approach-avoidance conflict on the predator-inhabited foraging task (the “robogator” task) showed behavioral changes, consistent both with previous studies and with a hypothesis that rats were showing increased concern about being attacked by the robot (worry). There was increased time spent hesitating in the nest, which led to a decrease in the number of laps run, and, hence, a decrease in the food reward received. Additionally, we observed an increase in the rate of midtrack aborts, in which rats ran out into the main track, but fled back to the nest before reaching the goal, and increased on-track pausing, in which rats paused on the track for a few seconds before proceeding.

Importantly, the hippocampal representations were fundamentally different during each of the three observed behaviors. During hesitation, the hippocampus was more likely to represent the robot and less likely to represent the food goal during high-synchrony events (HSEs, hippocampal cell bursts) after the rats had been attacked. Prior to engaging in a midtrack abort (MTA) there was a greater likelihood that the hippocampus was representing the attack threshold and robot, than in matched lap controls. The hippocampus remained in the theta state through the MTA event, and the hippocampal sweeps generally represented the rats’ planned direction of travel; however, after turning back towards the nest, late theta phases included representations of the robot and the attack threshold even though it was behind the animal. During on-track pausing, the non-local portion of theta cycles represented the robot and the far feeder, suggesting pauses arise from conflict between the danger to avoid and the goal to approach.

Transient representations of negatively-valued potential destinations during hippocampal SWRs have been seen on a novel shock-task [36], but this was on a novel maze, on which hippocampal representations during SWRs is known to be different [65,66], and it was not known what caused these hippocampal transient representations of the danger zone. As noted above, we also found increased representations of the robot during HSEs during hesitation in the nest. Fascinatingly, these observed increases in robot representation did not arise from changes in the sets of cells participating in the HSEs, but rather from a change in the spatial tuning of the hippocampal cells after the rat was attacked. When we used the post-attack tuning to decode hippocampal representations during HSEs in the nest, we found an increased representation of the robot, even with HSEs from before the attack. Conversely, there was no change in the robot representation during HSEs after the attack if we used the pre-attack tuning as the training-set for decoding. This remarkable result provides a fascinating mechanism for increased worry through hippocampal representations — random bursts of cell activity during HSEs and SWRs would pull an increased representation of the robot, not because there was attention to the robot during the HSEs and SWRs, but because the place cells had adjusted to include more robot-related fields.

One of the most interesting discoveries revealed in these data were the changes in spatial tuning after the first time the rat was attacked. As can be seen in Figures 3C and 3D, after being attacked during Session 5 (the first Attack session), the spatial tuning increased at the location of the robot and the area beyond the attack threshold. Importantly, however, these new fields were not due to the novelty of the robot itself, as the robot was present during the Novelty sessions (Sessions 3 and 4), but the spatial tuning did not change until the rat was attacked on Session 5.

Place fields have been observed to change after important episodic events, such as maze changes, attention-grabbing events such as sudden environmental manipulations, and the presence of novel objects (see [48] for review). Place fields have been seen to appear when novel objects are placed on the track [86–88], through direct induction of synaptic plasticity [89,90], or when rats stop and head-scan, presumably attending to distal external cues [73]. In fear-conditioning experiments, novel place fields have been seen to appear at the location of the animal after the first shock [71,72]. Importantly, however, all of these changes occurred at the location of the rat, presumably reflecting the rat’s personal contextual information at the time of the episodic event. Place field changes on the robogator task have been previously reported as being more likely to occur near the robot than near the nest [2,46], but the time course of those changes was unknown. Here, we observed the sudden appearance of new place fields and an increase in existing spatial activity, not at the location of the rat at the moment of attack, but rather at the location of the robot and space beyond the attack threshold — a new episodic memory of the object of worry, suggesting that the hippocampus can represent a novel event at a non-local location.

In contrast to hippocampal processing during hesitation in the nest, during midtrack aborts (MTAs), the hippocampus remained in the theta state. Consistent with approach tasks, the hippocampus represented where the animal was during the first half of theta, and future paths during the second half of theta [24,25,27,53–55]. During the MTA, just before the animal turned around, the representation in the second half of the theta cycle concentrated at the attack threshold, the point where the robot might attack the rat, making this the effective next choice point. As the animal turned, the representations in the second half of theta started to break apart, but contained increased information about that attack threshold, even as the animal started to flee back to the nest. Previous work has found that theta sweeps can proceed in a different direction from the orientation of the animal, but in these previous studies, theta sweeps have always been in the actual direction of motion [91,92]. These data suggest, instead, that cell firing in the second half of theta was representing attentional locations that the animal is concerned about, consistent with the transient jumps to the danger zone during the second half of theta seen on the active place avoidance task [56].

In contrast to both the hesitation on the nest and the midtrack aborts, the hippocampus remained in relatively normal theta during the on-track pauses, and the representations during the first and second halves of theta were relatively normal, although there was increased representation of the robot and feeder during the pause. It is likely that these on-track pauses were initiated by other decision-systems, not involving the hippocampus, but that information about the threat and reward was represented during the pause. To what extent, on-track pauses relate to freezing seen during active fear [18,93–96] or whether they relate to tonic immobility [97,98] remains unclear and will likely require further study.

Fascinatingly, diazepam affected all of these processes, in line with its effects on anxiety-like behavior. The effects of benzodiazepines on the hippocampus are relatively well studied, but not in vivo as animals are behaving on a decision-making or anxiogenic task. For instance, rodents allowed to move in an open field or walking on a forced-walk treadmill show a downward shift in theta frequency under diazepam [99–101], but it is not known whether this theta shift occurs during specific anxiety-related behaviors. Additionally, diazepam is known to interfere with sharp-wave ripple events [102–104], but it is not known whether these changes occur during anxiety-related behaviors.

We found that diazepam had the effect of increasing how long the rats spent in the nest, but, conversely, dramatically reduced the number of midtrack aborts and on-track behaviors. In line with these behavioral reductions, diazepam affected the underlying hippocampal processes associated with each of these. Diazepam reduced the central frequencies of theta and beta oscillations, and it obliterated the SWRs in the hippocampus. An interesting effect of diazepam was that it reversed theta asymmetry, which could dramatically affect ascending decoding.

Because the diazepam was delivered systemically, these effects could be due to primary effects of diazepam on hippocampal neurophysiology or to other systems impinging on hippocampus either directly or indirectly. However, these data raise the fascinating possibility that one mechanism through which diazepam achieves its anxiolytic effects is by interfering with hippocampal functionality and the ability to represent objects of worry.

Theories have long hypothesized that anxiety involves imagination, and worry involves negative episodic thinking [1–5]. Extensive data has identified a role for the hippocampus in the imagination of other places and other times [6,7,10,48], and, in particular in planning, particularly about positive futures to approach [9,24,25]. Here, we find that these same hippocampal processes also encode important information about negative events, both experienced dangers and potentially dangerous futures.

## Methods

### Subjects

A total of eleven (6 male, 5 female) Brown Norway rats aged between 7 and 10 months served as experimental subjects. Six (3 male, 3 female) rats were subjects in the neural ensemble recording experiment, and the remaining five (3 male, 2 female) rats were subjects in the diazepam experiment. All rats were maintained on a 14:10 hr light/dark cycle. During the experiment rats were food restricted such that their entire daily complement of food was earned from 45 mg food pellets (full-nutrition, Test Diet) in the foraging arena. Rats were always kept above 80% free-feeding weights and had unlimited access to water when in their home cages. All procedures were approved by the University of Minnesota (UMN) Institutional Animal Care and Use Committee (IACUC) and were performed in accordance with NIH guidelines.

The two cohorts of rats had to meet different criteria prior to the surgery. The first cohort of rats, which did not receive the drug manipulation, were trained to run a minimum of 50 laps on the linear track for two consecutive days. The second cohort was trained on the linear track for 7 days prior to the surgery. In both cases, the rats had ad libitum access to food (Teklad pellets) for at least 3 days prior to surgery.

### Surgery

Prior to the surgery we anesthetized the rat with 0.5-2% isoflurane mixed with medical-grade O_2_ in an induction chamber, and maintained this level of anesthesia throughout the surgery via a Somnosuite system (Kent Scientific, Torrington, CT). Once the rat was anesthetized, we placed the rat on a stereotax (Kopf Instruments, Tujunga CA). We removed the skull and dura above the dorsal hippocampus, chronically implanted silicon probes anterior-posterior at -3.8 and medial-lateral +2.5 or -2.5 (probe details in the next paragraph), affixed the probe and amplifier assembly by its connector to the skull surface with MetaBond (Parkell, Edgewood, NY), sealed the surgical hole with wax, and surrounded the assembly with a protective shroud that was printed on a 3D printer (Formlabs, Somerville, MA). After the surgery, the rats recovered in an incubator to maintain body temperature and were orally given 0.8 mL of Children’s Tylenol to alleviate discomfort. During the surgery and for the three days following the rats were administered 25 mg/kg of Baytril and 5 mg/kg of carprofen. During the post-surgery days we cleaned the surgical site with betadine. We resumed behavioral training with the rat after 3 days.

The silicon probes that we utilized varied by the experiment. With the first cohort of rats that served for the neural ensemble recordings (3 male, 3 female), we bilaterally implanted 64-channel, four shank Cambridge Neurotech P-1 probes into the dorsal hippocampus. With most of the diazepam cohort,we unilaterally implanted 64-channel, single-shankCambridge Neurotech H-3 silicon probes into the left hemisphere. One male rat in the diazepam cohort was unilaterally implanted into the left hemisphere with a 32-channel Cambridge, two-shank Neurotech E-2 silicon probe. The silicon probes used in the pharmacological experiment had been replated after being used in a prior experiment.

### Diazepam

A separate cohort of five rats were administered an anxiolytic drug manipulation of diazepam. Diazepam (2 mg/kg) was dissolved in Tween-20 to prepare a stock solution, which we then diluted with 0.9% saline. For our control manipulation we used a vehicle solution (10% Tween20 in saline). Both diazepam and Tween-20 were sourced from Sigma Adlrich (St. Louis MO). We chose this dose for diazepam due to its known efficacy in behaving rats [105] and previous behavioral experiments on this task [39]. We intraperitoneally administered all injections 5 min prior to each session. During the experiment we alternated the administration of diazepam with the vehicle solution within each subject.

### Statistical and Computing Resources

Data collection and analyses were conducted via a combination of MATLAB, Python, and R scripts. MATLAB (version R2015b; MathWorks, Natick, MA) was used to record behavioral data, manage the task, generate robot attacks, and deliver food. Python (version 3.9.4) code was used to perform data management, signal analysis, identification of behavioral events, and neural decoding. Notable Python packages that were utilized were *neuroDSP* (version 2.1.0), *numpy* (version 1.24.3), *scipy* (version 1.10.1), *pandas* (version 1.5.3), and *matplotlib* (version 3.6.2). R (version 4.2.2) was used to perform statistical analyses and plotting in RStudio. Notable R packages used were *car* (version 3.1-1), *dplyr* (version 1.0.10), *emmeans* (version 1.8.4-1), *forcats* (version 0.5.2), *ggplot2* (version 3.4.0), *lme4* (version 1.1-31), *performance* (version 0.10.2), and *tibble* (version 3.1.8).

### Statistical Analysis Techniques

Statistical analyses used a mixed-model framework by including a random effect of the subject within generalized linearized mixed effects regressions via the *lme4* R package. The number of laps, SWR rates, HSE rates were modeled as a Gaussian distribution. Since hesitation duration was non-Gaussian, we log-normalized it prior to fitting a linear mixed effects regression. HSE decodings were modeled as binomial distributions after coercing values less than 0.5 to 0 and values greater than 0.5 to 1. The likelihood of a midtrack abort on an outbound journey was also modeled as a binomial distribution. The number of pauses on a journey was modeled as a Poisson distribution. For significant interaction and main effects, we used the *emmeans* R package to determine significant differences. This analysis adjusted the degrees of freedom via the Kenward-Roger method and the p-values via the Tukey method.

To statistically assess changes in power spectral densities, we utilized dependent t-tests via the Python *scipy.stats.ttest_rel* function. Power of a frequency band was calculated from the average power across the frequency of its range. When we assessed the differences in spectral power at two time points, we compared the delta (1-4 Hz), theta (6-10 Hz), beta (15-20 Hz), and SWR (120-250 Hz) frequency bands. Since we were making multiple assessments across the PSD we used a Sidak correction to adjust statistical significance (alpha) values.

### Foraging Arena

The foraging arena was 111 cm in length and consisted of a nest, a track, and a robot bay on the side that formed an “L” shape that was constructed with DUPLO bricks (LEGO, Billund, Denmark). The DUPLO bricks were placed in a random and irregular color scheme to provide identifiable patterns to facilitate the formation of place cells. The nest was 19 cm long and 26 cm wide with partial walls creating a smaller opening that connected it to the track. The track was 67 cm long and 26 cm wide with the robot bay on the side at the far end from the nest. The robot bay was 22 cm along the side shared with the track and 35 cm deep. The robot was placed at an angle in the robot bay such that it would move towards the rat when it attacked. On sessions that the robot was not presented, a wall made of DUPLO bricks blocked off the robot bay.

Two feeders (Med-Associates, Georgia VT) were located on opposite sides of the track with one centered in the nest and one at the far end of the linear track. The latter feeder was on the opposite side of the robot, which meant that the rat had to pass by the robot to reach the delivered food pellet. The feeders provided food pellets when the rat approached it, but only if the rat had gone to the other feeder since the last visit. Two pellets were provided at each feeder to all rats for all sessions in the experiment that did not have a drug manipulation. The experiment with the drug manipulation had the feeders deliver two pellets to the female rats and three pellets to the male rats.

There were three session types across the experimental sessions: Linear Track, Novelty, and Attack. On Linear Track sessions the robot bay was walled off, which left the purely linear maze for the rat to run back and forth on. On Novelty sessions, the robot bay was open with the robot present but it did not engage in any actions during the session. On Attack sessions, the robot attack was triggered by the rat crossing an unmarked threshold. The first attack on these sessions could not occur until after the 15th lap. On this and subsequent laps, there was a 20% chance of the robot attacking when the rat crossed the threshold.

#### Robot

We used a robot constructed from the SPIK3R set (set number 31313, LEGO Mindstorms, Billund, Denmark), which has a scorpion-like shape and we modified the design for our purposes. The primary modification was that we removed the legs from that design and made it run on wheels along a grooved track. The robot was controlled by in-house programming (MATLAB), which used the position of the animal as a trigger to initiate an ‘attack’ behavior by the robot. When an attack was initiated the robot would make a screeching noise for a 1 second before surging forward and moving its arms. After the attack, the robot would move back to its starting position.

### Behavioral Quantification

We tracked the position of the rat by identifying the location of a red LED that was attached to the headstage. We tracked the LED by finding the position of maximum brightness within the confines of the foraging arena. If the position of maximum brightness was not above a value threshold, then we noted that we had lost track of the rat. The rat’s position was recorded at a rate of 30 Hz. From the change in the rat’s position, we identified a number of important behavioral events. When we normalized the data by subject, we subtracted the mean and divided by the standard deviation of that subject’s parameter values.

#### Hesitation

Hesitation was identified as anytime the rat was inside the nest or in the doorway to the nest. Hesitation time started when the rat entered the nest space from the track and ended when the rat left the doorway, proceeding out onto the track.

#### Mid-Track Aborts (MTAs)

We identified candidate MTAs as instances where the rat entered the track and returned to the door. As the minimal criteria, the rat’s headstage LED had to be on the track for 0.5 seconds and to have traveled 5 cm on the track. When we required a precise time for identification of the MTA itself, we visually assessed the candidate MTAs and removed instances when the animal did not at least turn their body to be perpendicular to the length of the track. Cases where a candidate MTA was identified but the animal did not turn typically indicated instances of the rat stretching onto the track without fully leaving the door zone, which resulted in them backing up into the nest. Given the qualitatively different nature of this action, we chose to exclude instances of it for MTA analysis. We identified the time of the MTA as when the rat’s head began to move in the direction of the eventual turn as this was the most consistent time point for analysis. Midtrack aborts were also identified when the animal was returning to the nest. In these cases we used the same criteria as the outbound journey, but did not go back to assess the exact time when engaging in the turn.

#### Pauses

We identified pause events from the behavioral tracking. Pauses were instances that the rat stopped moving for a prolonged period of time. Since the LED was attached to the rat’s headstage we ran a 2 Hz low pass filter to remove sharp movements. From this smoothed movement we identified instances when the animal moved by less than 1 cm along the long axis of the track for over half a second. The start and end of each pause were identified as when the animal began and stopped meeting this criteria, respectively.

### Cell Identification

Recordings were taken from using an Intan RhD 2000 system (Intan, Los Angeles CA) at 30 kHz. In the first experiment, a band-stop filter at 60 Hz was implemented at the time of recording. Voltage signals were first normalized by subtracting the median signal at each sample across the duration of the recording for common noise rejection. We then used Kilosort 2.0 [106] to sort the cellular activity into putative cells. We then used Phy [107] to visually assess the quality of the clusters and merge and split them as necessary. Following the spike-sorting process, we cross-correlated each putative cell’s spike waveforms with the median amplitude spike to determine the optimal temporal spike alignment and adjusted the spike timing appropriately. Putative pyramidal cells were identified as those cells with a mean firing rate less than 10 spikes/sec. We identified the locations of the probes post sacrifice by applying cresyl violet to coronal slices of the brain.

### Theta-Phase Identification

To differentiate the ascending and descending phases of theta we utilized a cycle-by-cycle technique [108]. We filtered the local field potential to specific and broad theta band frequencies of 6-10 Hz and 6-40 Hz, respectively. From the specific signal, we identified the simple peaks and troughs as the maxima and minima between 0 power crossings. From these points, we then looked at the broader local field potential, which better represents the saw-tooth pattern of theta in the hippocampus, and identified more accurate time points of the peak and trough.

### Power Spectral Density Analyses

For analyses of the local field potential’s power spectral density we used the spectrogram function from the scipy package, with a bin length of 1 second. When the power spectral density analysis needed to be aligned with an event (e.g., pauses and midtrack aborts) we used the spectrogram function to define custom time bins for the analysis around the event. Given that these events were typically briefer we used a bin length of 0.25 seconds. After computing the spectrograms, we used Welch’s method to determine the power spectral density for the separately identified before and after event time periods. When comparing power spectral densities we defined the frequency bins using standard frequencies. Delta was defined as 1-4 Hz, theta as 6-10 Hz, beta as 15-20 Hz, gamma as 30-80 Hz. Sharp-wave ripples were defined as the more narrow 150-250 for identifying individual events, but a broader 120-250 Hz for power spectral density analysis. We used the wider band for many analyses because we observed that the cross frequency correlations that typically indicate the sharp-wave frequency band began at that point with the rats we were working with (see, for example, Figure 4A). To compare power spectral densities, we used paired and unpaired t-tests of the frequency band powers, and corrected for the multiple comparisons via the Šidák method [109].

### Sharp-Wave Ripple Channel Identification

To identify sharp-wave ripples, we found the channel that had the greatest oscillatory power in 120-250 Hz range above and beyond the aperiodic activity. We identified this channel for each probe shank by calculating the power spectral density (PSD) of each channel on the shank, removing the aperiodic component, and then comparing the residual power within the power spectrum. We identified the aperiodic component by fitting a robust linear regression to the log-log representation of the PSD in the 80-400 Hz frequency range. This approach was taken because the log-log representation of the PSD linearizes the aperiodic and we used a robust linear regression so that the influence of the SWR power on the regression was minimized. After identifying the aperiodic component, we removed it from the PSD that gave us the remainder, which was primarily the SWR power. The best candidate SWR layer was identified as the channel on the shank with the largest SWR power in the 120-250 Hz frequency range. We qualitatively validated the best SWR layer by visually examining the LFPs after bandpass filtering it between 120 and 250 Hz, ensuring that the identified channel had good SWRs.

### Sharp Wave Ripple (SWR) Identification

Sharp wave ripple (SWR) events were defined as instances when the local field potential power in the 150-250 Hz range was more than four standard deviations above its mean for more than 20 ms. To determine when this criteria was met, we bandpass filtered the local field potential and applied a Hilbert transform to shift it into the complex number domain. We normalized the power and then identified candidate SWR events by determining when the power was above the criteria threshold. We merged candidate instances when the gap between the end and start was less than 5 ms or when the gap between the start of two instances was less than 20 ms. After this merge, we filtered these candidates to just those instances that lasted longer than 20 ms.

### High-Synchrony Event (HSE) Identification

We identified high-synchrony events (HSE) by identifying bursts of pyramidal cell activity. We summed the spiking of putative pyramidal cells in 1 ms bins. Next, we applied a Gaussian kernel with a standard deviation of 7 ms to these bins. From this data, we identified candidate HSEs as those instances when the kerneled spike activity was more than three standard deviations above the mean over the session. From this set of candidates, we only kept instances during which the higher spike activity was maintained for more than 20 ms and less than 750 ms with the starts and ends of these instances being when they started and stopped meeting the greater than three standard deviations criterion.

### Changes in spatial tuning

When examining the changes in spatial tuning we segmented the track into 64 bins along the length of the track and 3 bins for the direction of travel (moving towards the robot, stationary, moving towards the nest). When constructing cellular tuning curves on these dimensions, we only used cellular activity while the hippocampal LFP was dominated by theta and only during its descending phase. This approach should minimize the influence of cellular activity during sharp wave ripples and theta forward sweeps, and thus maximize the local spatial accuracy. From the cellular activity during these periods we constructed the tuning curve, and then normalized it by the rats’ occupancy in the space while traveling in the specified direction. For statistical comparisons, we collapsed over the length of spatial bins that comprised each zone.

### Decoding

We used a naive Bayesian approach [110] to decode the spatial information represented by ensemble neural activity in the dorsal hippocampus. For the decodings, we segmented constructed tuning curves using the same method as described previously. Ascending and descending theta phase were decoded separately (as a whole, taking into account the variable time of each phase). High synchrony events were decoded as a single whole, decoded over the duration of the event.

## Author contributions

OLC designed the experiment, performed the experiment that recorded cellular activity, conducted all analyses reported here, and wrote and edited the primary manuscript. MTE performed the diazepam experiment and worked with ADR to do an initial analysis. CJW designed the initial experiment, which served as the basis for this project. ADR helped design the experiment, helped with analyses, wrote and edited the paper and supervised the project.

## Acknowledgements and funding

This work was funded by R01-MH080318 (ADR), a T32 fellowship to OLC (T32-DA037183), a T32 fellowship to CJW (T32-DA030672), summer project funding from St. Olaf College to MTE, and funding from the University of Minnesota Medical School. We are indebted to Chris Boldt, Kelsey Seeland, and Ayaka Sheehan for technical work on this project as well as Kevin Singh for help with running animals. We thank Chelsey Damphousse, Geoff Diehl, Jonathan Gewirtz, Shmuel Lissek, and Ugurcan Mugan for comments on earlier versions of this work.

